# ATF4 Expression in Thermogenic Adipocytes is Required for Cold-Induced Thermogenesis in Mice via FGF21-Independent Mechanisms

**DOI:** 10.1101/2023.03.09.531964

**Authors:** Sarah H. Bjorkman, Alex Marti, Jayashree Jena, Luis M Garcia Pena, Eric T. Weatherford, Kevin Kato, Jivan Koneru, Jason Chen, Ayushi Sood, Matthew J. Potthoff, Christopher M. Adams, E. Dale Abel, Renata O. Pereira

**Author notes:** **Lead contact:**, Address: 169 Newton Road, 4338 PBDB | Iowa City, IA 52242. Sarah H. Bjorkman and Alex Marti should be considered joined first-authors.

## Abstract

In brown adipose tissue (BAT), short-term cold exposure induces the activating transcription factor 4 (ATF4), and its downstream target fibroblast growth factor 21 (FGF21). Induction of ATF4 in BAT in response to mitochondrial stress is required for thermoregulation, partially via upregulation of FGF21. In the present study, we tested the hypothesis that *Atf4* and *Fgf21* induction in BAT are both required for BAT thermogenesis by generating mice selectively lacking either *Atf4 (*ATF4 BKO) or *Fgf21* (FGF21 BKO) in UCP1-expressing adipocytes. After 3 days of cold exposure, core body temperature was significantly reduced in *ad-libitum*-fed ATF4 BKO mice, which correlated with *Fgf21* downregulation in brown and beige adipocytes, and impaired browning of white adipose tissue (WAT). Conversely, despite having reduced browning, FGF21 BKO mice had preserved core body temperature after cold exposure. Mechanistically, ATF4, but not FGF21, regulates amino acid import and metabolism in response to cold, likely contributing to BAT thermogenic capacity under *ad libitum*-fed conditions. Importantly, under fasting conditions, both ATF4 and FGF21 were required for thermogenesis in cold-exposed mice. Thus, ATF4 regulates BAT thermogenesis by activating amino acid metabolism in BAT in a FGF21-independent manner.

**Graphical Abstract:** 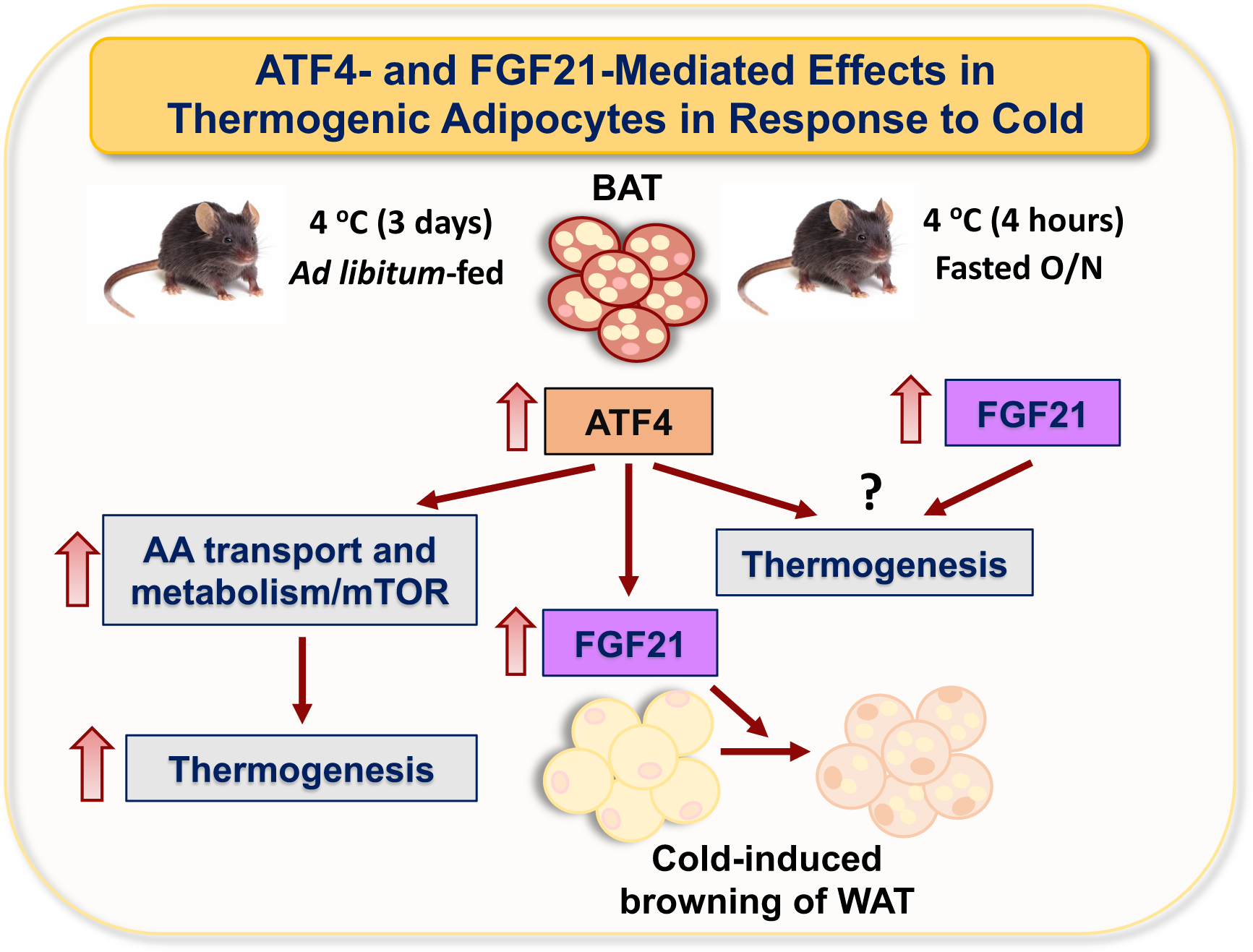

## Introduction

Since the discovery of active brown adipose tissue (BAT) in adult humans, multiple studies have explored BAT thermogenic activation as a potential strategy for increasing energy expenditure and nutrient disposal to mitigate obesity and its complications (1–3). More recently, BAT has been recognized as a secretory organ, promoting the release of batokines that may act in an endocrine manner to regulate systemic metabolism (4). Indeed, several studies using β3-adrenergic agonists to activate BAT demonstrated changes in adiposity and improvements in glucose homeostasis (3, 5, 6), however, this class of medication poses cardiovascular concerns (1). Therefore, identifying new pathways that can be targeted to induce BAT’s thermogenic activity and secretory function, may lead to the discovery of new targets to improve metabolic health.

A recent study demonstrated that the activating transcription factor 4 (ATF4), the main effector of the integrated stress response (ISR), is induced in brown adipocytes in response to cold exposure, which correlates with a significant increase in fibroblast growth factor 21 (FGF21) circulating levels (7). Furthermore, ATF4 overexpression in BAT is sufficient to improve thermogenesis in mice (8). Similarly, our recent work showed that selective deletion of the mitochondrial fusion protein optic atrophy 1 (OPA1) in BAT induces the expression and secretion of fibroblast growth factor 21 (FGF21) as a batokine, via an ATF4-dependent mechanism. Activation of this ATF4-FGF21 axis was required to mediate an adaptive response characterized by elevated resting metabolic rates, and improved thermoregulation in OPA1-deficient mice (9). Together, these studies suggest that the ISR is likely to be operative in BAT under physiological circumstances and may regulate the secretion of FGF21 as a batokine. However, studies investigating the ISR in BAT are few and were performed in whole-body knock out models and cultured adipocytes (10–14), limiting our understanding of the specific roles of ATF4 in BAT. Moreover, although FGF21 is induced in BAT in response to thermogenic activation, whether ATF4 is required for this induction, and whether BAT-derived FGF21 is necessary for adaptive thermogenesis was yet to be investigated.

In the present study, we investigated the requirement of ATF4 and FGF21 induction in thermogenic adipocytes for adaptive thermogenesis by generating mice with selective deletion of either *Atf4* or *Fgf21* in UCP1-expressing adipocytes. Our data demonstrated that neither ATF4 nor FGF21 are required for the activation of the thermogenic gene program in BAT after 3 days of cold-exposure, however, *Atf4* expression in BAT was required for the cold-induced upregulation of *Fgf21* and both *Atf4* and *Ffg21* expression was required for cold-induced browning of white adipose tissue (WAT). Nonetheless, core body temperature was only significantly reduced in ATF4 BKO mice, but not in FGF21 BKO mice, when mice were cold exposed with free access to food. Noteworthy, in the absence food, both *Fgf21* and *Atf4* were required for thermoregulation in mice. Together, our study identified novel roles for ATF4 in BAT physiology in regulating core body temperature in a manner that is independent of *Fgf21* induction in thermogenic adipocytes or cold-induced browning. Our data points to an alternative mechanism downstream of *Atf4*, which involve regulation of amino acid transport and metabolism to support thermogenesis.

## Research Design and Methods

### Animals and Animal Care

Experiments were performed in male mice on a C57Bl/6J background, unless otherwise stated. ATF4^fl/fl^ (15) and FGF21^fl/fl^ mice (16) were generated as previously described. Transgenic mice expressing cre recombinase under the control of the *Ucp1* promoter (Tg (Ucp1-cre)1Evdr) (17) were acquired from the Jackson Laboratories (#024670), and were crossed with ATF4^fl/fl^ and FGF21^fl/fl^ mice to promote selective deletion of these genes in thermogenic adipocytes (ATF4 BKO and FGF21 BKO mice, respectively). ATF4^fl/fl^ and FGF21^fl/fl^ mice not expressing the Cre recombinase were used as wild type (WT) controls.

Mice were weaned at 3 weeks of age and kept on standard chow (2029X Harlan Teklad, Indianapolis, IN, USA). Animals were housed at 22 ^°^C with a 12-h light, 12-h dark cycle with free access to water and standard chow, unless otherwise noted. For cold exposure experiments, mice were acclimated to 30 ^°^C (thermoneutral temperature for mice) for 7 days to dampen brown adipose tissue thermogenesis prior to being cold-exposed (4 ^°^C). All mouse experiments presented in this study were conducted in accordance with animal research guidelines from the National Institutes of Health (NIH) and were approved by the University of Iowa Institutional Animal Care and Use Committee (IACUC).

### Cold exposure experiments

Core body temperature telemeters (Respironics, G2 E-Mitter, Murrysville, PA, USA) were surgically implanted into the abdominal cavity of 8-10-week-old mice. The mice were then allowed to recover for 6 days post-surgery, while individually housed in a rodent environmental chamber (Power Scientific, Pipersville, PA, USA) at 30 °C. Mice were then transferred to an OxyMax Comprehensive Lab Animal Monitoring System (CLAMS, Columbus Instruments International) at 30 °C for 3 days, followed by 4 °C for 3 days while singly housed, as previously described (18). Core body temperature was recorded every 17 minutes throughout the experiment, along with O_2_ and CO_2_ levels, food intake, and ambulatory activity, as estimated by photoelectric beam breaks in the X + Y plane. Mice were fed *ad libitum* throughout the study. A separate cohort of mice underwent acute cold exposure following fasting. For these studies, 12-week-old mice were initially individually housed in the rodent environmental chamber at 30 °C for 7 days. At the end of the seventh day, mice were fasted for 12 hours (7 pm – 7 am). The initial temperature (t0) was recorded using a rectal probe (Fisher Scientific, Lenexa, KS, USA) at 7 am on day 8, the temperature was switched to 4 °C. Once the desired temperature was reached, we recorded rectal temperatures hourly for up to 4 hours during cold exposure. Mice remained without food for the entirety of the experiment.

### Glucose tolerance tests, nuclear magnetic resonance, and serum analysis

Glucose tolerance tests (GTT) and measurements of fasting glucose levels were performed as previously described (9). Serum FGF21 (BioVendor ELISA kit, Asheville, NC, USA) and GDF15 (R&D Systems, Minneapolis, MN, USA) were measured using commercially available kits according to the manufacturers’ directions. Whole body composition was measured by nuclear magnetic resonance in the Bruker Minispec NF-50 instrument (Bruker, Billerica, MA, USA) (9).

### Analysis of triglyceride levels

Triglycerides levels were measured in the liver of ATF4 BKO mice, using the EnzyChrom™ Triglyceride Assay Kit (BioAssay Systems, Hayward, CA, USA), as previously described (9).

### RNA extraction and quantitative RT–PCR

Total RNA was extracted from tissues with TRIzol reagent (Invitrogen) and purified with the RNeasy kit (Qiagen Inc, Germantown, MD, USA), as previously described (19). Additional details and primer sequences can be found in Supplementary Material.

### RNA Sequencing

RNA sequencing was performed in BAT of male ATF4 BAT KO mice cold exposed for 3 days by the Iowa Institute of Human Genetics: Genomics Division at the University of Iowa. Additional details on animal studies can be found in Supplementary Material.

### Western blot analysis

Immunoblotting analysis was performed as previously described (20). Approximately, 50 mg of frozen tissue was homogenized in 200 μl lysis buffer containing (in mmol/l) 50 HEPES, 150 NaCl, 10% glycerol, 1% Triton X-100, 1.5 MgCl_2_, 1 EGTA, 10 sodium pyrophosphate, 100 sodium fluoride, and 100 μmol/l sodium vanadate. Right before use, HALT protease/phosphatase inhibitors (Thermo Fisher Scientific, Waltham, MA, USA) were added to the lysis buffer and samples were processed using the TissueLyser II (Qiagen Inc., Germantown, MD, USA). Tissue lysates were resolved on SDS–PAGE and transferred to nitrocellulose membranes (Millipore Corp., Billerica, MA, USA). Membranes were incubated with primary antibodies overnight at 4 °C and with secondary antibodies for 1 h, at room temperature. The data was analyzed using Image Studio Lite (LI-COR Technologies, Lincoln. NE, USA) and was normalized by the specified loading controls. Data is represented as arbitrary units of optical density (OD). An antibody list can be found in Supplementary Material.

### Data analysis

Unless otherwise noted, all data are reported as mean ± SEM. Student’s *t*-test was performed for comparison of two groups. ANOVA followed by the appropriate post-hoc analysis was performed for comparisons of 3 or more groups. A probability value of *P* ≤ 0.05 was considered significantly different. Statistical calculations were performed using the GraphPad Prism software (La Jolla, CA, USA). The CalR software was used to extract the averaged data for the metabolic parameters measured in the CLAMS at 30 °C and 4 °C for both, the light and dark cycles. The hourly plots for body temperature were also extracted from CalR (21).

## Results

### ATF4 and its downstream targets are induced in BAT after 3 days of cold exposure

ATF4 was shown to be potently induced in BAT in response to acute cold-stress (4-6 hours) following a 12-hour fast, which correlated with high serum levels of FGF21 and growth differentiation factor 15 (GDF15), downstream targets of ATF4 (7). We therefore, first sought to examine if this pathway was also induced after 3 days of cold exposure in *ad libitum*-fed conditions. Mice were acclimated to thermoneutrality (30 °C) for 7 days to attenuate brown adipose tissue thermogenic function, after which they were exposed to 4 °C for 3 days to induce thermogenesis. Our data demonstrated increased phosphorylation of the eukaryotic translation initiation factor 2A (eIF2α), which is upstream of ATF4 induction (Figure 1A). Accordingly, mRNA levels of *Atf4*, and its downstream targets *Fgf21* and *Gdf15* were also induced in BAT (Figure 1B) in response to 3-days of cold exposure. To investigate whether ATF4 induction in BAT is required for adaptive thermogenesis, we generated mice lacking ATF4 selectively in thermogenic adipocytes (ATF4 BKO mice). *Atf4* expression was significantly decreased in BAT of KO mice (Figure 1C), while *Atf4* expression was preserved in the inguinal white adipose tissue (iWAT) (Figure 1D) at room temperature conditions. *Fgf21* and *Gdf15* mRNA (Figure 1E) expression in BAT and their respective circulating levels (Figure 1F and G) were unchanged between WT and KO mice under baseline conditions (chow-fed mice at room temperature).

**Figure 1:**
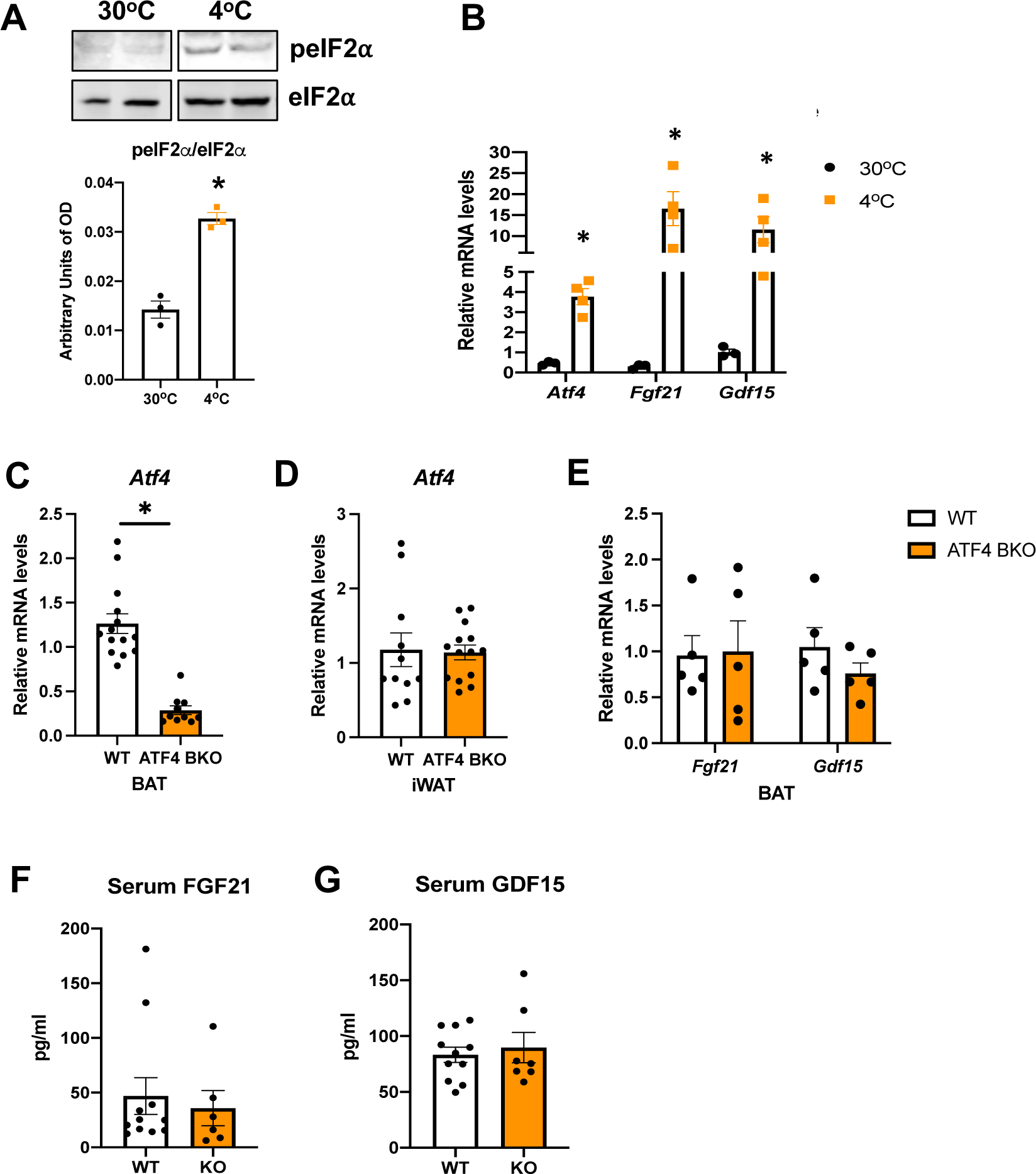
ATF4 and its downstream targets are induced in BAT after 3 days of cold exposure. **A and B.** Data collected in BAT of WT mice exposed to 30 ^°^C or 4 ^°^C for 3 days. **A.** Immunoblot of phosphorylated eIf2α normalized to total eIf2α and respective densitometric quantification. Images cropped from the same blot. Optical density (OD). **B.** mRNA expression of *Atf4*, *Fgf21* and *Gdf15.* **C-G.** Data collected in WT and ATF4 BAT KO male and female mice at baseline conditions. **C.** Relative mRNA expression of *Atf4* in BAT normalized to *Gapdh.* **D.** Relative mRNA expression of *Atf4* in inguinal white adipose tissue (iWAT) normalized to *Gapdh.* **E.** Relative mRNA expression of *Fgf21* and *Gdf15* in BAT normalized to *Gapdh.* ISR (integrated stress response.) **F.** Serum FGF21 levels. **G.** Serum GDF15 levels. Data are expressed as means ± SEM. Significant differences were determined by Student’s *t*-test, using a significance level of *P* < 0.05. * p < 0.05 significantly different vs. WT mice or 30 ^°^C.

### ATF4 BAT KO mice have reduced BAT mass and modestly impaired glucose tolerance at baseline conditions

Because global ATF4 deletion has been associated with changes in systemic metabolism (12, 13), we evaluated body composition, and glucose homeostasis in 8-week-old ATF4 BKO mice. Body mass (Fig. 2A) and total lean mass (Fig. 2B) were unchanged between KO mice and their WT littermate controls, while total fat mass was increased in KO mice (Fig. 2C). Although total fat mass was increased, brown adipose tissue (BAT) mass was modestly, but significantly reduced in ATF4 BKO mice (Fig. 2D). Interestingly, UCP1 protein levels were also significantly reduced in BAT of ATF4 BKO mice at baseline conditions (Fig. 2E), while rectal core body temperature was unchanged between genotypes (Fig. 2F). To assess the impact of *Atf4* deletion in thermogenic adipocytes on glucose homeostasis, we performed glucose tolerance tests (GTT). ATF4 BKO mice had a modest impairment in glucose tolerance (Fig. 2J), as demonstrated by increased area under the curve for the GTT (Fig. 2H), and higher fasting glucose levels after a 6-hour fast (Fig. 2I), while liver triglyceride levels were comparable between genotypes (Fig. 2J). Hence, under baseline conditions, ATF4 deletion in BAT has minor effects in systemic metabolic health. We believe that the mild impairments in glucose homeostasis is likely a consequence of the small increase in fat mass observed in these animals.

**Figure 2:**
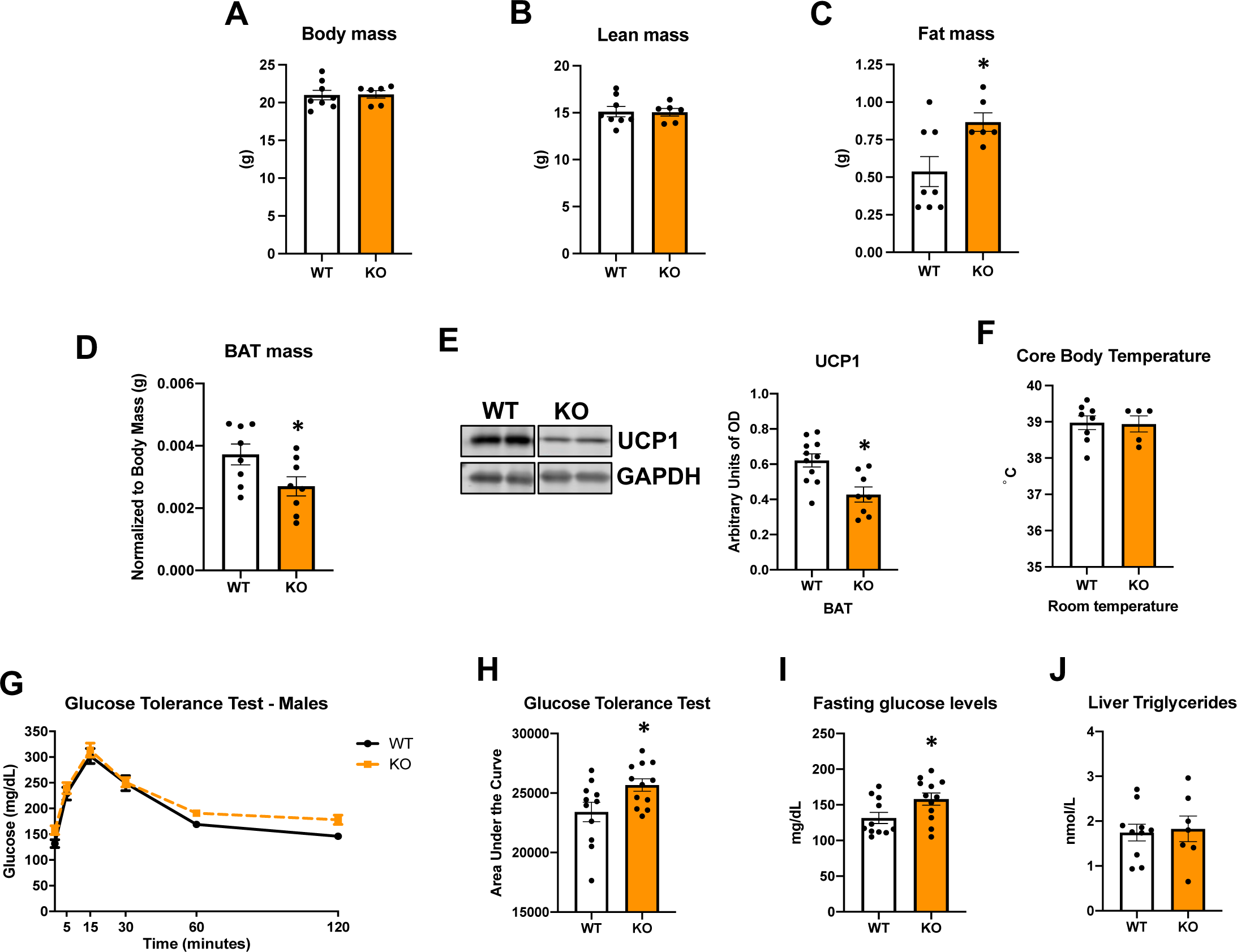
ATF4 BAT KO mice have reduced BAT mass and modestly impaired glucose homeostasis at baseline conditions. Data collected under baseline conditions in ATF4 BKO (KO) male mice and their respective WT littermate controls. **A.** Body mass. **B.** Total lean mass. **C.** Total fat mass. **D.** Brown adipose tissue (BAT) mass normalized to body mass. **E.** UCP1 protein levels in BAT normalized to GAPDH and respective densitometric quantification. Images cropped from the same blot. Optical density (OD). **F.** Core body temperature measured with a rectal probe. **G.** Glucose tolerance test (GTT). **H.** Area under the curve for the GTT. **I.** Fasting glucose levels (after a 6-hr fast). **J.** Liver triglyceride levels. Data are expressed as means ± SEM. Significant differences were determined by Student’s *t*-test, using a significance level of *P* < 0.05. * p < 0.05 significantly different vs. WT mice.

### ATF4 expression in BAT is required for thermoregulation in mice

To determine the requirement of ATF4 expression in BAT for adaptive thermogenesis, we monitored core body temperature and metabolic changes in ATF4 BKO mice and their WT littermate controls during 3 days at thermoneutrality (30 °C) and 3 days of cold exposure (4 °C). Although averaged core body temperature was unchanged between WT and KO mice at 30 °C, core body temperature was significantly reduced in KO mice during cold exposure (Fig. 3A and B). Despite having reduced thermoregulatory capacity, the metabolic phenotype of ATF4 BKO mice was largely unaffected, with no significant changes in energy expenditure (Fig. 3C), food intake (Fig. 3D), or locomotor activity (Fig. 3E) at thermoneutrality or during cold exposure, when compared to WT littermate controls (no effect associated with genotype). Mice were euthanized and tissues were harvested after the 3 days of cold exposure for molecular analysis. Cold-induced mRNA expression of *Fgf21* (Fig. 3F) was significantly reduced in ATF4 BKO mice, but serum levels were unaffected by *Atf4* deficiency in BAT (Fig. 3G). Reduced core body temperature correlated with a small, but significant decrease in BAT mass (Fig. 3H), however, relative mRNA expression of thermogenic genes was unchanged between genotypes (Fig. 3I). UCP1 protein levels (Fig. 3J) were modestly decreased in BAT of KO mice after cold exposure. Browning of WAT is induced in response to 3 days of cold exposure, and is believed to contribute to thermoregulation in mice (18). We, therefore, assessed browning of iWAT in WT and ATF4 BKO mice after 3 days of cold exposure. *Atf4* mRNA expression was significantly reduced in iWAT of KO mice after cold exposure (Fig. 3K). Moreover, activation of the thermogenic gene program was attenuated (Fig. 3L), and UCP1 protein levels were significantly reduced in iWAT of KO mice (Fig. 3M), suggesting impaired cold-induced browning. Together these data suggest that ATF4 expression in thermogenic adipocytes is required to support cold-induced thermogenesis in mice, and cold-induced browning in iWAT. Furthermore, ATF4 also regulates *Fgf21* mRNA expression in thermogenic adipocytes in response to cold. However, whether FGF21 induction is thermogenic adipocytes is required to mediate thermogenesis remained to be determined.

**Figure 3:**
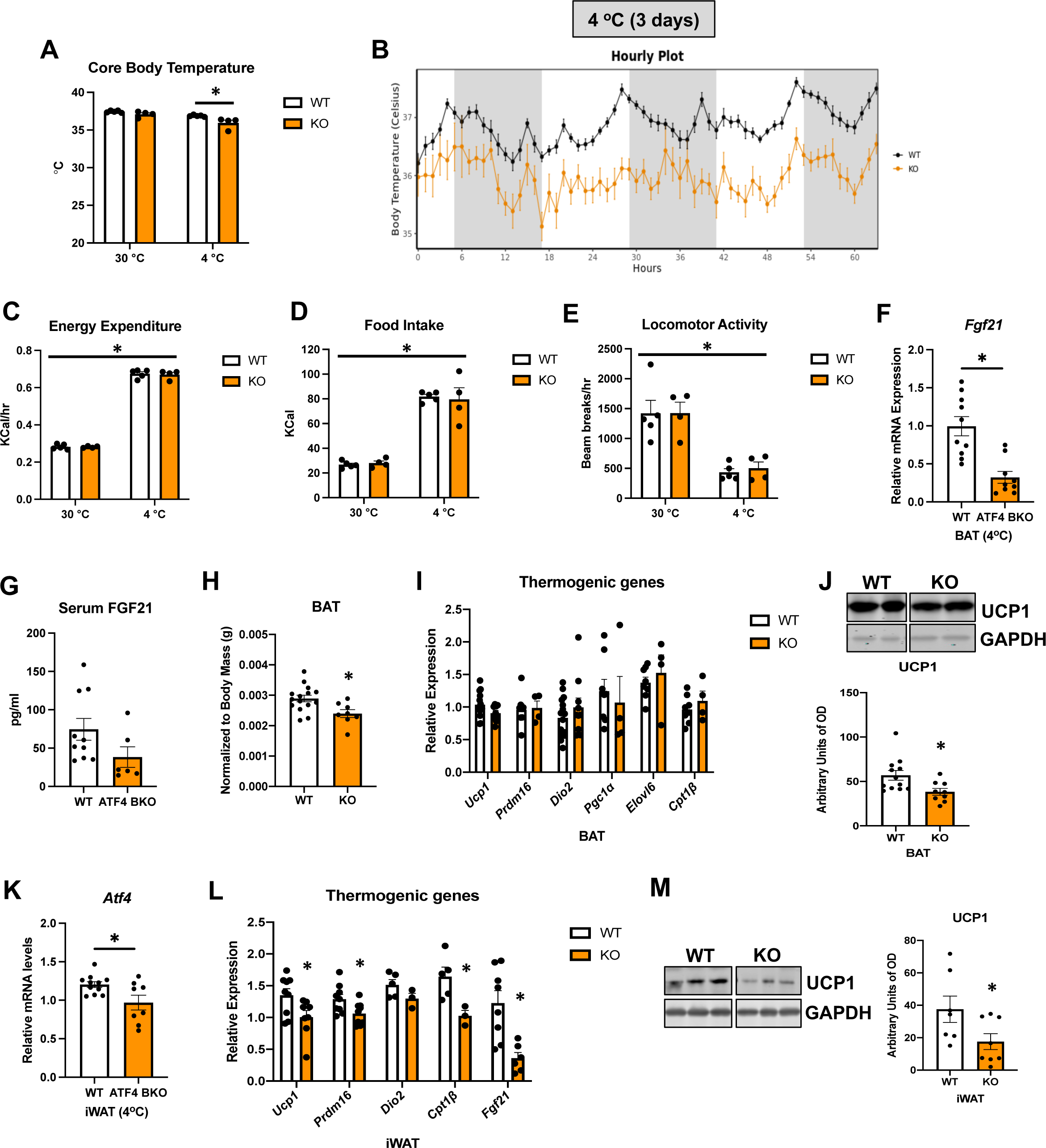
ATF4 expression in BAT is required for thermoregulation in mice. **A-E.** Data collected in WT and ATF4 BKO (KO) male mice at either during 3 days at thermoneutrality (30 ^°^C) or during 3 days of cold exposure (4 ^°^C). **A.** Averaged core body temperature. **B.** Core body temperature over time during cold exposure. **C.** Energy expenditure. **D.** Food intake. **E.** Locomotor activity. **F-M.** Data collected in WT and ATF4 BKO (KO) male mice after 3 days of cold exposure (4 ^°^C). **F**. Relative *Fgf21* mRNA expression in BAT normalized to *Gapdh*. **G.** Serum FGF21 levels. **H.** BAT mass normalized to body mass. **I.** mRNA expression of thermogenic genes in BAT normalized to *Gapdh*. **J.** Immunoblot of UCP1 in BAT normalized to GAPDH and its respective densitometric quantification. Images cropped from the same blot. Optical density (OD). **K.** mRNA expression of *Atf4* in inguinal white adipose tissue (iWAT) normalized to *Gapdh*. **L.** mRNA expression of thermogenic genes in iWAT normalized to *Gapdh*. **M.** Immunoblot of UCP1 in inguinal white adipose tissue (iWAT) normalized to GAPDH and its respective densitometric quantification. Images cropped from the same blot. Optical density (OD). Data are expressed as means ± SEM. Significant differences were determined by Student’s *t*-test, using a significance level of p < 0.05. * Significantly different vs. WT mice or 30 ^°^C.

### FGF21 BAT KO mice have normal body weight and glucose homeostasis at baseline conditions

To test the role of *Fgf21* expression in thermogenic adipocytes for adaptive thermogenesis, we selectively deleted *Fgf21* in *Ucp1*-expressing adipocytes (FGF21 BKO). Because global FGF21 deletion has been associated with changes in glucose and energy homeostasis (16, 22), we evaluated body composition, and glucose homeostasis in 8-week-old FGF21 BKO mice fed chow diet. Body mass (Fig. 4A), total lean mass (Fig. 4B) and total fat mass (Fig. 4C) were unchanged between KO mice and their WT littermate controls. BAT weight normalized to body weight (Fig. 4D), UCP1 protein levels in BAT (Fig. 4E) and core body temperature (Fig. 4F) were also similar between WT and KO mice under baseline conditions. To assess the impact of FGF21 deletion in thermogenic adipocytes on glucose homeostasis, we performed glucose tolerance tests (GTT). FGF21 BKO mice had similar glucose clearance as WT mice, as demonstrated by unchanged areas under the curve for the GTT (Fig. 4G and H), and comparable fasting glucose levels after a 6-hour fast (Fig. 4I).

**Figure 4:**
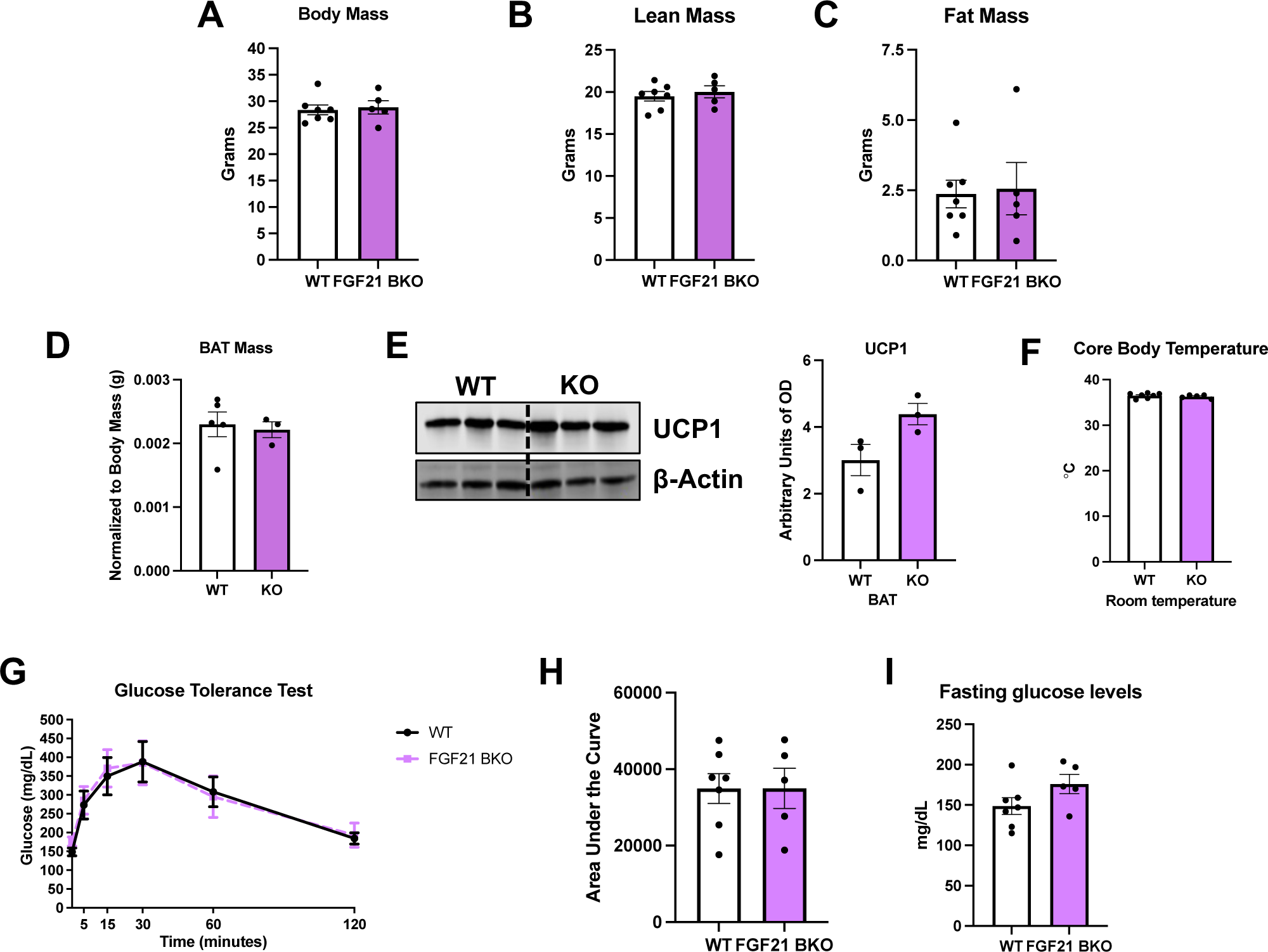
FGF21 BAT KO mice have normal body weight and glucose homeostasis at baseline conditions. Data collected under baseline conditions in FGF21 BKO (KO) male mice and their respective WT littermate controls. **A.** Body mass. **B.** Total lean mass. **C.** Total fat mass. **D.** Brown adipose tissue (BAT) mass normalized to body mass. **E.** UCP1 protein levels in BAT normalized to β-Actin and respective densitometric quantification. Optical density (OD). **F.** Core body temperature measured with a rectal probe. **G.** Glucose tolerance test (GTT). **H.** Area under the curve for the GTT. **I.** Fasting glucose levels (after a 6-hr fast). Data are expressed as means ± SEM. Significant differences were determined by Student’s *t*-test, using a significance level of *P* < 0.05.

### FGF21 expression in thermogenic adipocyte is dispensable for thermoregulation

*Fgf21* mRNA expression was significantly reduced in ATF4 BKO mice after cold exposure. Therefore, we directly tested the role of cold-induced *Fgf21* expression in thermogenic adipocytes for adaptive thermogenesis by exposing FGF21 BKO mice to 4 °C for 3 days. We monitored core body temperature and metabolic changes in FGF21 BKO mice and their WT littermate controls for 3 days at thermoneutrality (30 °C) and 3 days of cold exposure (4 °C). Averaged core body temperature, as measured by telemetry, was unchanged between WT and FGF21 BKO mice at thermoneutrality (Fig. 5A) or after 3 days of cold exposure (Fig. 5A and B). Of note, 1 WT mouse and 2 KO mice died unexpectedly soon after the temperature was changed to 4 °C and were removed from the analysis. Regarding energy homeostasis, energy expenditure (Fig. 5C) and locomotor activity (Fig. 5E) were unaffected in KO mice either at 30 °C or 4 °C. Food intake was significantly increased in WT mice after cold exposure, but not in KO mice (Fig. 5D). Although *Fgf21* mRNA levels were significantly reduced in BAT (Fig. 5F), FGF21 serum levels (Fig. 5G) were unchanged between genotypes, indicating that BAT-derived FGF21 does not contribute to FGF21 circulating levels after 3 days of cold exposure. BAT mass normalized to body weight was unaffected by FGF21 deletion (Fig. 5H). Likewise, expression of thermogenic genes (Fig. 5I), and UCP1 protein levels were similar in BAT of WT and FGF21 BKO mice (Fig. 5J). Next, we evaluated the effects of FGF21 deletion on cold-induced browning of iWAT. *Fgf21* mRNA levels were significantly reduced in iWAT of cold-exposed FGF21 BKO mice (Fig. 5K), which correlated with reduced mRNA levels of the thermogenic genes *Ucp1* and *Ppargc1α* (Fig. 5L), and with significantly lower UCP1 protein levels (Fig. 5M). Together, our data reinforce a role for FGF21 on cold-induced browning of WAT and suggest that impaired browning in the absence of defective BAT thermogenesis is insufficient to hamper thermoregulation in mice, at least under the conditions tested in this study. Our data also indicate that defective *Fgf21* expression in ATF4 BKO mice is likely dispensable for the impaired thermoregulatory capacity observed in these mice. Finally, our data suggest that FGF21 induction in BAT does not contribute to FGF21 serum levels in response to cold exposure, and it is dispensable for cold-induced thermogenesis in *ad libitum*-fed mice.

**Figure 5:**
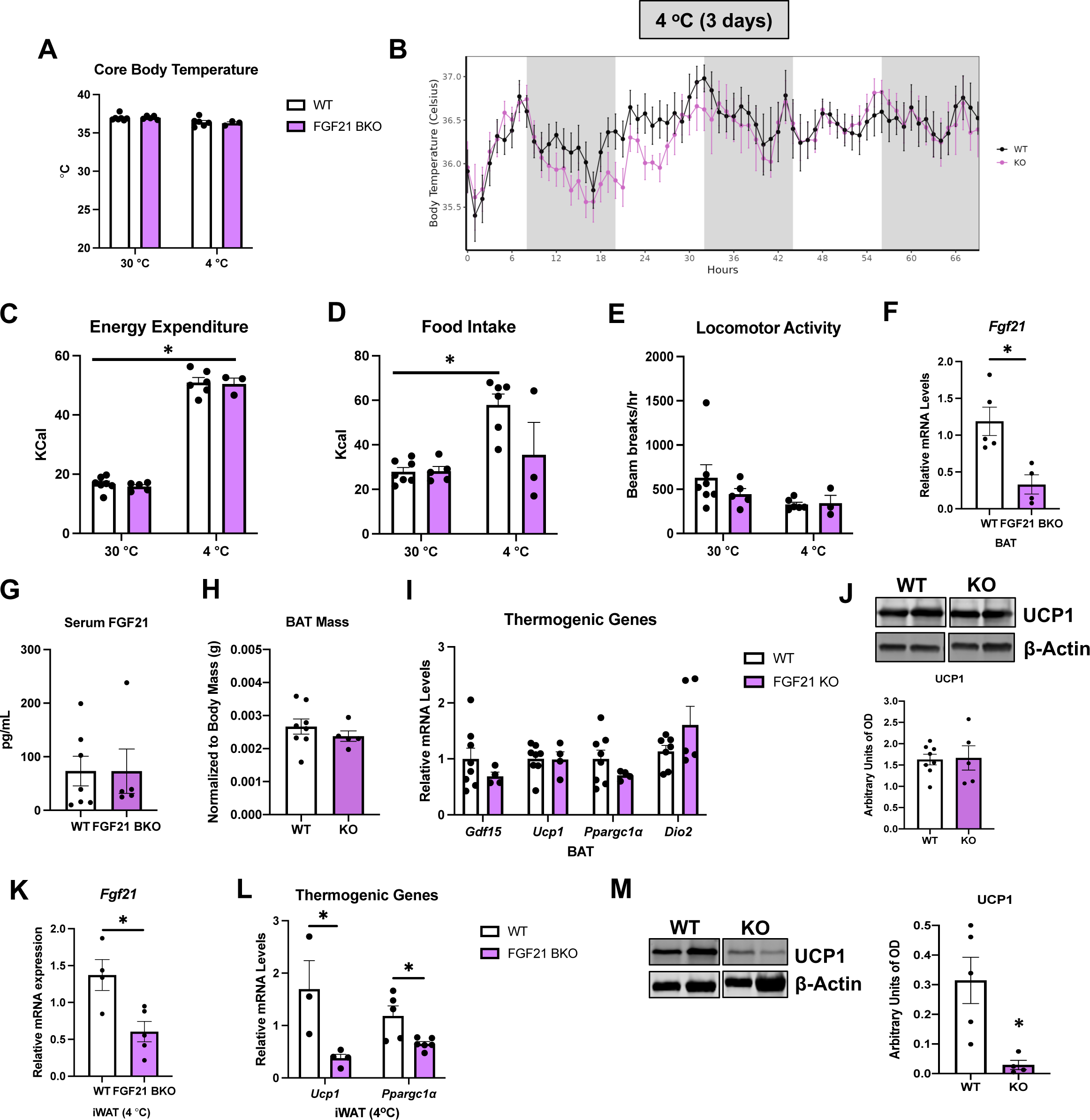
FGF21 Expression in Thermogenic Adipocytes Is Dispensable for Thermoregulation. **A-E.** Data collected in WT and FGF21 BKO (KO) male mice at either during 3 days at thermoneutrality (30 ^°^C) or during 3 days of cold exposure (4 ^°^C). **A.** Averaged core body temperature. **B.** Core body temperature over time during cold exposure. **C.** Energy expenditure. **D.** Food intake. **E.** Locomotor activity. **F-M.** Data collected in WT and FGF21 BKO (KO) male mice after 3 days of cold exposure (4 ^°^C). **F**. Relative *Fgf21* mRNA expression in BAT normalized to *Tbp* (tata box protein). **G.** Serum FGF21 levels. **H.** BAT mass normalized to body mass. **I.** mRNA expression of thermogenic genes in BAT normalized to *Tbp*. **J.** Immunoblot of UCP1 in BAT normalized to β-Actin and its respective densitometric quantification. Images cropped from the same blot. Optical density (OD). **K.** mRNA expression of *Fgf21* in inguinal white adipose tissue (iWAT) normalized to *Tbp*. **L.** mRNA expression of thermogenic genes in iWAT normalized to *Tbp*. **M.** Immunoblot of UCP1 in inguinal white adipose tissue (iWAT) normalized to β-Actin and its respective densitometric quantification. Images cropped from the same blot. Optical density (OD). Data are expressed as means ± SEM. Significant differences were determined by Student’s *t*-test, using a significance level of p < 0.05. * Significantly different vs. WT mice or 30 ^°^C.

### Transcriptome analysis reveals downregulation of genes involved in amino acid metabolism in ATF4 BKO mice

To gain insight into the molecular mechanisms driving impaired thermoregulation in ATF4 BKO mice, we performed RNA sequencing (RNASeq) in BAT collected from 10-12-week-old mice lacking ATF4 in thermogenic adipocytes (OPA1 BAT KO) and their respective wild-type (WT) littermate controls after 3 days of cold exposure. In addition to confirming downregulation of *Fgf21* in ATF4 BKO mice, our transcriptome data revealed that several amino acid transporters were amongst the top downregulated genes in ATF4-deficient BAT (Fig. 6A). Furthermore, Ingenuity Pathway Analysis uncovered that genes involved in amino acid synthesis and metabolism were repressed in cold-exposed ATF4 BKO mice relative to their WT counterparts (Fig. 6B). Repression of amino acid metabolism-related genes in ATF4 BKO mice after 3 days of cold exposure was confirmed by qPCR (Fig. 6C); Meanwhile, these genes were unchanged in BAT of cold-exposed FGF21 BAT KO mice under the same conditions (Supplemental Fig. 1). Of note, defective brain chain amino acids (BCAA) uptake and oxidation in BAT leads to reduced thermogenesis in mice (23), suggesting that BCAA catabolism in BAT is required for proper thermoregulation. Furthermore, a recent study demonstrated that ATF4 overexpression in BAT improves cold-induced thermogenesis, which seems to be dependent on increased mTOR signaling activation (8). Our data shows that mTOR signaling pathway, as measured by phosphorylated ribosomal protein S6, was reduced in BAT of cold-exposed ATF4 BKO mice relative to WT controls (Figure 6D). Together, these data suggest that ATF4, but not FGF21 is required to regulate amino acid import and metabolism into BAT during cold exposure, and may also regulate mTOR signaling in response to cold, thereby contributing to thermogenic regulation in mice.

**Figure 6:**
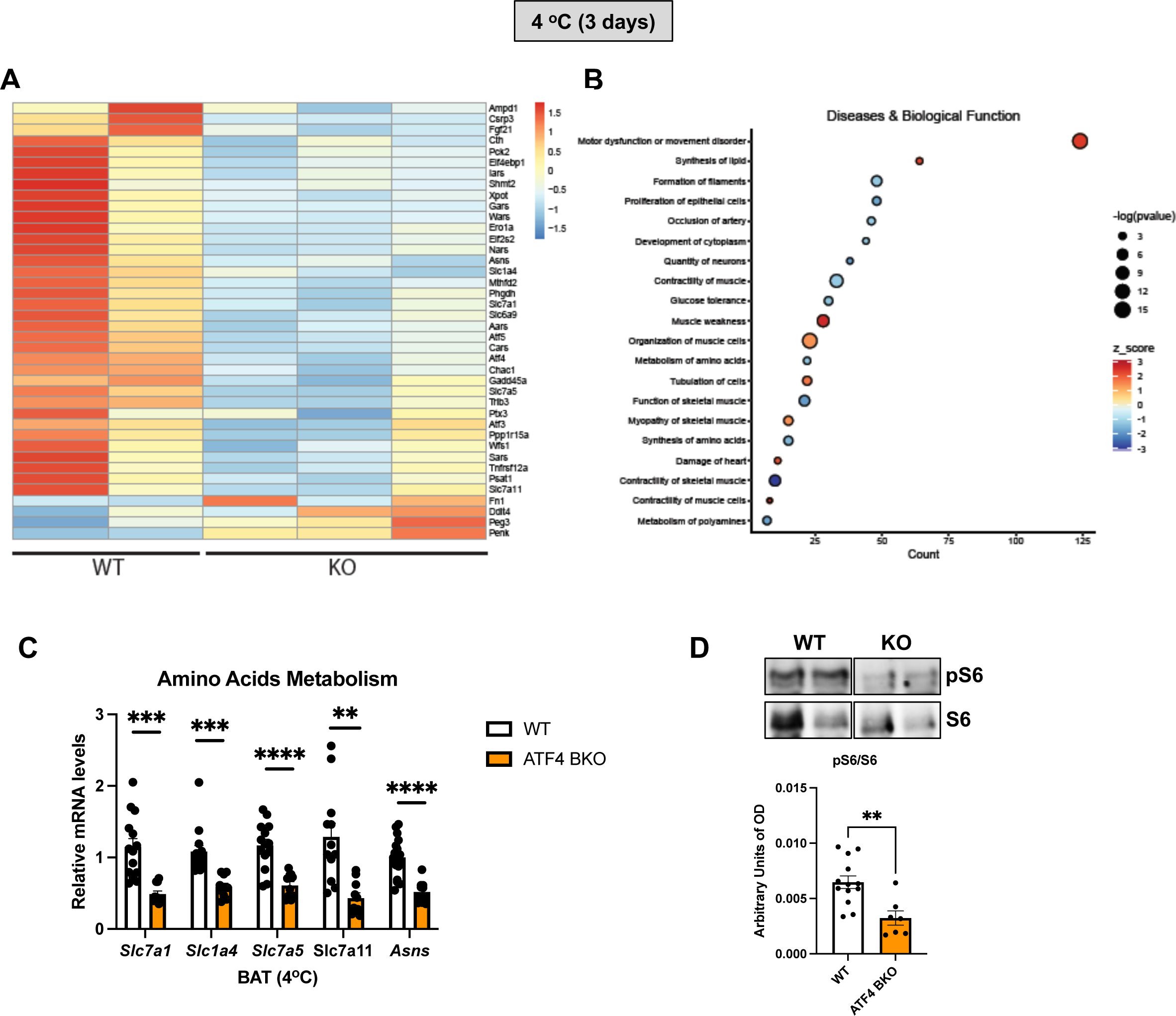
Transcriptome Analysis Reveals Downregulation of Genes Involved in Amino Acid Metabolism in ATF4 BKO Mice. **A-B.** Data collected from brown adipose tissue (BAT) of WT and ATF4 BKO mice cold exposed for 3 days (Differential gene expression and Ingenuity Pathway Analysis). **A.** Heatmap of differentially expressed genes annotated in the IPA database as ATF4 targets. Plotted are log2(x+1) transformed counts scaled by row. **B.** Genes differentially expressed have a significant overlap (adjusted p-value ≤ 0.05) with genes that are annotated to be involved in various diseases or have common biological functions that regulate genes with a significant overlap (adjusted p-value ≤ 0.05) to those differentially expressed in ATF4KO BAT. The number of genes in the data set annotated as being involved in the disease or have a particular biological function is plotted on the x-axis. Bubble size = -log (adjusted p-value) and color = z-score; indicates the potential impact of the differential gene expression pattern on the disease or biological function. **C.** Relative mRNA expression of amino acid transporter and metabolism genes in BAT of ATF4 BKO mice normalized to *Tbp* expression. **D.** Immunoblot of phosphorylated S6 in BAT normalized to total S6 and its respective densitometric quantification. Images cropped from the same blot. Optical density (OD). Data are expressed as means ± SEM. Significant differences were determined by Student’s *t*-test, using a significance level of p<0.05. ** p < 0.01; *** p < 0.001; **** p < 0.0001 significantly different vs. WT mice.

### ATF4 and FGF21 expression in thermogenic adipocytes is required to maintain core body temperature in mice cold exposed under fasting conditions

An earlier study reporting potent activation of ATF4 in BAT in response to acute cold exposure was performed following fasting, which correlated with an exacerbated induction in FGF21 serum levels (7). To test the role of ATF4 in thermoregulation and in FGF21 induction in BAT under these conditions, a separate cohort of ATF4 BKO mice was cold-exposed for 4 hours following a 12-hour fast. Core body temperature was significantly reduced in ATF4 BKO relative to their WT counterparts (Fig. 7A). Interestingly, our data also demonstrated that ATF4 is not required to regulate FGF21 expression in BAT (Fig. 7B) or FGF21 serum levels (Fig. 7C) in mice during acute cold exposure following fasting. Indeed, FGF21 secretion from liver is likely the primary contributor to FGF21 serum levels during fasting (24). Surprisingly, a subset of thermogenic genes was significantly induced in BAT of ATF4 BKO mice (Fig. 7D), while UCP1 protein levels were unchanged between ATF4 BKO mice and WT control mice in BAT under these conditions (Fig. 7E), suggesting normal UCP1-depdendent thermogenesis. Moreover, activation of thermogenic genes in iWAT was similar between WT and ATF4 BKO mice (Fig. 7F). The expression of AA transporters was unchanged in BAT of ATF4 BKO (Supplemental Fig. 2), suggesting ATF4 activates different pathways during cold exposure depending on food availability. Although FGF21 BKO mice were able to properly thermoregulate when cold exposed under *ad libitum*-fed conditions, in the absence of food they had reduced tolerance to cold relative to their WT counterparts (Fig. 7G). *Fgf21* mRNA levels were significantly reduced in BAT (Fig. 7H), but FGF21 serum levels (Fig. 7I) were similar between genotypes. Thermogenic gene expression (Fig. 7J) and UCP1 protein levels (Fig. 7K) in BAT were also unchanged between FGF21 BKO mice and their WT controls. Interestingly, cold-induced activation of thermogenic genes in iWAT was reduced in FGF21 BKO mice under fasting conditions (Fig. 7L). Together, our data suggest ATF4 and FGF21 expression in BAT are independently required to regulate core body temperature in mice during acute cold exposure following fasting.

**Figure 7:**
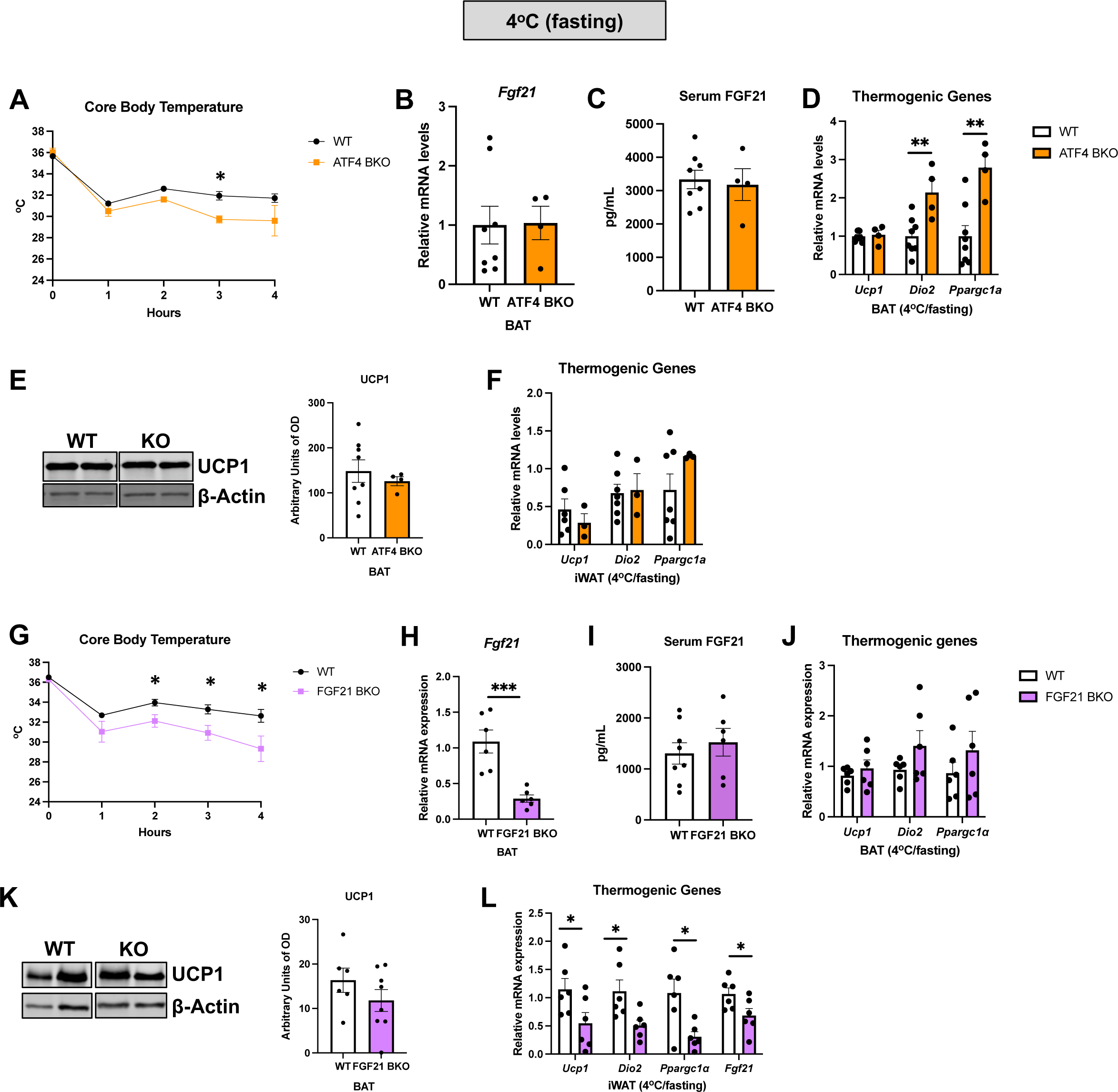
ATF4 and FGF21 Expression in Thermogenic Adipocytes Is Required to Maintain Core Body Temperature in Mice Cold Exposed Under Fasting Conditions. **A-F.** Data collected in WT and ATF4 BKO mice cold exposed for 4 hours following a 12-hr fast. **A.** Hourly core body temperature. B. *Fgf21* mRNA levels in BAT of ATF4 BKO mice normalized to *Tbp.* C. FGF21 serum levels. **D.** mRNA expression of thermogenic genes in BAT normalized to *Tbp*. **E.** Immunoblot of UCP1 in BAT normalized to β-actin and its respective densitometric quantification. Images cropped from the same blot. Optical density (OD). **F.** mRNA expression of thermogenic genes in iWAT normalized to *Tbp*. **G-L.** Data collected in WT and FGF21 BKO mice cold exposed for 4 hours following a 12-hr fast. **G.** Hourly core body temperature. H. *Fgf21* mRNA levels in normalized to *Tbp.* I. FGF21 serum levels. **J.** mRNA expression of thermogenic genes in BAT normalized to *Tbp*. **K.** Immunoblot of UCP1 in BAT normalized to β-actin and its respective densitometric quantification. Images cropped from the same blot. Optical density (OD). **L.** mRNA expression of thermogenic genes in iWAT normalized to *Tbp*. Data are expressed as means ± SEM. Significant differences were determined by Student’s *t*-test, using a significance level of p < 0.05. * p < 0.05; ** p < 0.01; *** p < 0.001 significantly different vs. WT mice.

## Discussion

ATF4 was recently shown to be potently induced in BAT in response to acute cold exposure, which correlated with increased FGF21 serum levels (7). Intriguingly, mice with global deletion of the *Atf4* gene have increased energy expenditure and induced *Ucp1* expression in BAT (13). Furthermore, these mice are better able to maintain core body temperature during a 7-h cold challenge (14). Nonetheless, a recent study demonstrated that adult-onset conditional *Atf4* deletion in agouti-related peptide neurons in the hypothalamus of mice is sufficient to increase energy expenditure, and enhance thermogenesis in BAT, suggesting that this phenotype is likely centrally regulated, rather than a result of *Atf4* deficiency in BAT (25). Indeed, our recently published study demonstrated that deletion of the mitochondrial protein optic atrophy 1 (OPA1) in BAT leads to upregulation of *Atf4*, which is required for the induction and secretion of FGF21 as a batokine and to improve thermoregulation in mice. Mice with concomitant deletion of OPA1 and either ATF4 or FGF21 specifically in thermogenic adipocytes had impaired thermoregulatory capacity in response to 3 days of cold exposure (4 °C) (9). Moreover, a recent study showed that selective activation of ATF4 in brown adipocytes improves cold-induced thermogenesis in mice (8). Therefore, we hypothesized that activation of ATF4 in response to cold is required for proper thermoregulation in mice and to induce FGF21 as a batokine.

Thermogenic stimulation has been shown to strongly induce *Fgf21* expression in BAT via mechanisms that involve canonical activation of β-adrenergic signaling pathways (18). However, in our recent study, we demonstrated that FGF21 can be induced even when BAT’s thermogenic function is dampened, via ATF4 activation (9). ATF4 binds to the FGF21 promoter and induces its transcription in several cell models (26–28). However, whether ATF4 is required for cold-induced upregulation of FGF21 in BAT remained to be elucidated. In the present study, we show that mice lacking ATF4 specifically in thermogenic adipocytes (ATF4 BKO) have reduced core body temperature after 3 days of cold exposure in the absence of major changes in the expression of thermogenic genes in BAT. Additionally, energy expenditure was unchanged in these animals at thermoneutrality and during cold exposure. Although thermogenic gene activation was maintained in BAT, cold-induced *Fgf21* mRNA expression was reduced in thermogenic adipocytes (BAT and iWAT), and cold-induced browning of WAT was attenuated in ATF4 BKO mice. Our data demonstrate that ATF4 induction is required for *Fgf21* upregulation in response to short-term cold stress in BAT and iWAT. Therefore, next we directly tested whether FGF21 is required to promote cold-induced thermogenesis, by generating mice lacking FGF21 selectively in thermogenic adipocytes.

Cold exposure is known to stimulate FGF21 expression in both humans and animal models; however, the source of circulating FGF21 and its metabolic consequences for adaptive thermogenesis are still incompletely understood (29, 30). A study in FGF21 global KO mice showed that ablation of FGF21 leads to an impaired response to cold stress when placing mice from 27 °C to 5 °C for 3 days (18). In UCP1 knockout (KO) mice, BAT was suggested as a potential source of circulating FGF21 upon long-term cold exposure, which was associated with increased browning of subcutaneous WAT relative to WT mice. However, a recent study in UCP1-FGF21 double knockout mice demonstrated that FGF21 is dispensable for this phenotype, as well as for the changes in energy expenditure and thermoregulation during prolonged cold exposure (31). Furthermore, a recent study demonstrated that liver-derived, but not adipose-derived FGF21 is required to maintain core body temperature in the first few hours of cold exposure (32). Therefore, although FGF21 may be dispensable for the adaptation to gradual long-term cold exposure in UCP1 KO mice, liver-derived FGF21 is likely required for thermoregulation in more acute scenarios. Although we observed no changes in FGF21 serum levels after 3 days of cold exposure, our data in ATF4 BKO mice, in which cold-induced FGF21 expression is diminished in BAT and WAT, could suggest that FGF21 expression in thermogenic adipocytes might be necessary for the regulation of core body temperature in response to short-term cold exposure in mice, maybe via autocrine/paracrine actions. However, our data in FGF21 BKO mice revealed no changes in BAT thermogenic activation or core body temperature after 3 days of cold exposure, even though cold-induced browning of WAT was attenuated. These data reinforce a role for FGF21 on cold-induced browning of WAT, as previously reported (18, 33, 34), and suggest that impaired browning is insufficient to hamper effective thermoregulation in mice when BAT thermogenesis is preserved. Our data also indicate that defective *Fgf21* expression in ATF4 BKO mice is unlikely to play a role on the impaired thermoregulatory capacity observed in these mice.

Transcriptome data in BAT of cold-exposed ATF4 BKO mice revealed that, in addition to reduced *Fgf21* transcript levels, gene expression of several amino acid transporters, including of brain chain amino acids (BCAA), and amino acid metabolism genes was repressed relative to WT control mice. A recent study showed that cold stimuli potently increase mitochondrial BCAA uptake and oxidation in BAT, leading to enhanced BCAA clearance in the circulation. Furthermore, defective BCAA catabolism in BAT leads to impaired BCAA clearance and reduced thermogenesis in mice (23), suggesting that BCAA catabolism in BAT is required to support thermogenesis. Furthermore, over the last several years, the mechanistic target of rapamycin (mTOR) signaling pathway has been reported to affect BAT function in rodents. Studies from independent groups showed that mTORC1 activity is highly induced by acute and chronic cold exposure and β3-adrenergic receptor stimulation (35–37). Furthermore, loss of mTORC1 in adipocytes reduces BAT size and completely prevents cold-induced BAT expansion, mitochondrial biogenesis, and oxidative metabolism in mice (36–38). Here, we show that phosphorylation of the downstream mTOR target ribosomal protein 6 (S6) is reduced in ATF4-deficient BAT after 3 days of cold exposure, suggesting that ATF4 induction in response to cold may regulate mTOR pathway to promote thermogenesis in BAT. Indeed, a recent study showed that ATF4 overexpression in BAT improves cold-induced thermogenesis, which seems to be dependent on increased S6 phosphorylation, and is attenuated upon treatment with the mTOR inhibitor rapamycin (8).

Of note, in the present study we reveal a novel role for FGF21 expression in thermogenic adipocytes in the modulation of core body temperature under fasting conditions. Although FGF21 BKO mice showed no change in core body temperature when cold exposed under *ad libitum*-fed conditions, acute cold exposure following an overnight fast reduced core body temperature in FGF21 BKO mice to a similar degree as observed in ATF4 BKO mice. Under these conditions, ATF4 was dispensable to regulate *Fgf21* levels in BAT, and FGF21 serum levels were similar in both ATF4 and FGF21 BKO mice relative to their WT counterparts, suggesting that liver is the predominant source of FGF21 under these conditions (24, 32). Although additional studies are needed to clarify the molecular mechanisms underlying these changes in core body temperature, here we show that they occur independently of changes in UCP1 protein levels in BAT of ATF4 BKO or FGF21 BKO mice. Furthermore, although browning of iWAT was unaffected in ATF4 BKO mice, FGF21 BKO had reduced expression of thermogenic genes in iWAT, suggesting attenuated browning.

In conclusion, by using mouse models of selective deletion of *Atf4* and *Fgf21* in thermogenic adipocytes, our study unraveled new roles for ATF4 and FGF21 for thermoregulation in mice. Together, our data demonstrated that *Atf4* deletion in thermogenic adipocytes impairs thermoregulation in mice, in a manner that seems independent of ATF4-mediated *Fgf21* expression or browning of WAT. Rather, ATF4 BKO mice had reduced expression of amino acid transporter genes and impaired mTOR signaling, suggesting ATF4 might contribute to BAT thermogenesis by increasing amino acid metabolism and by activating mTOR signaling in response to cold exposure. In contrast, FGF21 BKO mice had preserved thermoregulation, and normal activation of thermogenic genes in BAT, despite having reduced browning of WAT. Importantly, in the absence of food, both ATF4 and FGF21 are independently required to maintain core body temperature in mice regardless of changes in UCP1 levels.

## Acknowledgment

Metabolic phenotyping was performed at the Metabolic Phenotyping Core at the Fraternal Order of Eagles Diabetes Research Center. Analysis of mRNA expression was performed at the Genomics Division at The Iowa Institute of Human Genetics.

## Author Contributions

S.H.B. and A.M. designed and conducted the experiments, analyzed data and wrote the manuscript. E.T.W., L.M.G.P, J.J. and K.K., conducted experiments and analyzed data. K.K., J.K., J.C. and A.S. aided with animal work and processed tissues for biochemical analysis. M.J.P., C.A. and E.D.A. provided essential materials and critical expertise. R.O.P. conceived the project, coordinated all aspects of this work, and helped with manuscript writing and editing. E.D.A. assisted with manuscript editing.

