## Supplemental Files for "ATF4 Expression in Thermogenic Adipocytes is Required for Cold-Induced Thermogenesis in Mice via FGF21-Independent Mechanisms"

**Supplemental Experimental Procedures and Figure Legends**

Sarah H. Bjorkman[1,2*]; Alex Marti[1*]; Jayashree Jena[1]; Luis M Garcia Pena[1]; Eric T. Weatherford[1]; Kevin Kato[1], Jivan Koneru[1], Jason Chen[1], Ayushi Sood[1], Matthew J. Potthoff[1,3]; Christopher M. Adams[1,4]; E. Dale Abel[1,5]; Renata O. Pereira[1]

[1]Fraternal Order of Eagles Diabetes Research Center and Division of Endocrinology and Metabolism, Roy J. and Lucille A. Carver College of Medicine, University of Iowa, Iowa City, IA, USA.

[2]Department of Obstetrics and Gynecology, Reproductive Endocrinology and Infertility, University of Iowa Hospital and Clinics, Iowa City, IA, USA

[3]Department of Neuroscience and Pharmacology, Roy J. and Lucille A. Carver College of Medicine, University of Iowa, Iowa City, IA, USA.

[4]Division of Endocrinology, Diabetes, Metabolism and Nutrition, Department of Medicine, Mayo Clinic, Rochester, Minnesota, USA.

[5]Department of Medicine, David Geffen School of Medicine, University of California, Los Angeles, Los Angeles, CA, USA

Address: 169 Newton Road, 4338 PBDB | Iowa City, IA 52242

**Supplementary Methods:**

**RNA extraction and quantitative RT–PCR**

RNA concentration was determined by measuring the absorbance at 260 and 280 nm using a spectrophotometer (NanoDrop 1000, NanoDrop products, Wilmington, DE, USA). Total RNA (1 µg) was reverse-transcribed using the High-Capacity cDNA Reverse Transcription Kit (Applied Biosystems, Waltham, MA, USA), followed by qPCR reactions using SYBR Green (Life Technologies, Carlsbad, CA, USA) (1). Samples were loaded in a 384-well plate in triplicate, and real-time polymerase chain reaction was performed with an ABI Prism 7900HT instrument (Applied Biosystems, Waltham, MA, USA). The following cycle profile was used: 1 cycle at 95°C for 10 min; 40 cycles of 95°C for 15 s; 59°C for 15 s, 72°C for 30 s, and 78°C for 10 s; 1 cycle of 95°C for 15 s; 1 cycle of 60°C for 15 s; and 1 cycle of 95°C for 15 s. Data were calculated using the delta-delta Ct method and normalized to either *Gapdh* or *Tbp* expression. Results are shown as relative mRNA levels. qPCR primers were designed using Primer-Blast or previously published sequences (2). Utilized primers are listed in Table 1.

| **Gene name** | **Forward** | **Reverse** |
|---|---|---|
| *Fgf21* | TGACGACCAAGACACTGAAGC | TTTGAGCTCCAGGAGACTTTCTG |
| *Atf4* | AGCAAAACAAGACAGCAGCC | ACTCTCTTCTTCCCCCTTGC |
| *Ucp1* | GTGAAGGTCAGAATGCAAGC | AGGGCCCCCTTCATGAGGTC |
| *Prdm16* | CAGCACGGTGAAGCCATTC | GCGTGCATCCGCTTGTG |
| *Gapdh* | AACGACCCCTTCATTGAC | TCCACGACATACTCAGCAC |
| *Dio2* | AATTATGCCTCGGAGAAGACCG | GGCAGTTGCCTAGTGAAAGGT |
| *Cpt1b* | TGCCTTTACATCGTCTCCAA | AGACCCCGTAGCCATCATC |
| *Evlov6* | TCAGCAAAGCACCCGAAC | AGCGACCATGTCTTTGTAGGAG |

| Gene name | Forward | Reverse |
|---|---|---|
| *Ppargc1$\alpha$* | GTAAATCTGCGGGATGATGG | AGCAGGGTCAAAATCGTCTG |
| *Gdf15* | GAGAGGACTCGAACTCAGAAC | GACCCCAATCTCACCTCTG |
| *Slc7a1* | CGT GAG TAC GCG ATC CTT GT | AGG ACC AAG ATG GAC TCG GA |
| *Slc1a4* | GTG GCA TCG CTG TTG CTT AC | GAC GTA GTG AAT GCG GCA AC |
| *Slc7a11* | GTC TGC CTG TGG AGT ACT GT | ATT ACG AGC AGT TCC ACC CA |
| *Slc7a5* | ATC GTA GGT CCT GCC ATG TG | ACC GTG TCT GAG CTA GTT GC |
| *Asns* | TAC AAC CAC AAG GCG CTA CA | AAG GGC CTG ACT CCA TAG GT |
| *Tbp* | ACC CTT CAC CAA TGA CTC CTA TG | TGA CTG CAG CAA ATC GCT TGG |

**Supplemental Table 1. Primer sequences**

**RNA Sequencing**

Sequencing libraries were prepared using the Illumina TruSeq mRNA Stranded kit and sequenced on a HiSeq4000. Two approaches were used for read alignment, mapping, and quantification. First, a workflow using HISAT2 (v2.1.0), featureCounts (v1.6.3), and DESeq2 (v1.22.2) was performed (3-5). The second approach used pseudo-alignment and quantification with Kallisto (v0.45.0) and DESeq2 for differential expression analysis (6). Ingenuity® Pathway Analysis (IPA®) software from Qiagen was utilized for identification of potentially modified pathways. Data visualization was performed using pheatmap and ggplot2 packages in R. RNA-seq data have been deposited to the GEO database under the accession number GSE227060.

**Immunoblotting and antibody list:**

Approximately, 50 mg of frozen tissue was homogenized in 200 µl lysis buffer containing (in mmol/l) 50 HEPES, 150 NaCl, 10% glycerol, 1% Triton X-100, 1.5 $MgCl_2$, 1 EGTA, 10 sodium pyrophosphate, 100 sodium fluoride, and 100 µmol/l sodium vanadate. Right before use, HALT protease/phosphatase inhibitors (Thermo Fisher Scientific, Waltham, MA, USA) were added to the lysis buffer and samples were processed using the TissueLyser II (Qiagen Inc., Germantown, MD, USA). Tissue lysates were resolved on SDS–PAGE and transferred to nitrocellulose membranes (Millipore Corp., Billerica, MA, USA). Membranes were incubated with primary antibodies overnight at 4 °C and with secondary antibodies for 1 h, at room temperature. The data was analyzed using Image Studio Lite (LI-COR Biotechnology, Lincoln, NE, USA) and was normalized by the specified loading controls. Data is represented as arbitrary units of OD.

**Antibodies**

Primary Antibodies: GAPDH (1:1,000, Cell Signaling Technology, Danvers, MA, USA, #2118), UCP1 (1:1,000, Abcam, Boston, MA, USA, #Ab10983), phosphorylated eIF2α serine 51 (1:1,000, Cell Signaling Technology, #3597), total eIF2α (1:500, Santa Cruz Biotechnology, Dallas, TX, USA, #SC81261), phosphorylated S6 (1:1,000, Cell Signaling Technology, #4858), total S6 (1:1,000, Cell Signaling Technology, #2317) and β-actin (1:1,000, Sigma Aldrich, St. Louis, MO, USA, # A2066). Secondary antibodies: IRDye 800CW anti-mouse (1:10,000, LI-COR, Lincoln, NE, USA, #925-32212) and Alexa Fluor anti-rabbit 680 (1:10,000, ThermoFisher Scientific, #A27042). Fluorescence was quantified using the LiCor Odyssey imager.

**Supplemental Figure Legends:**

**Supplemental Figure 1:** mRNA expression of amino acid metabolism genes normalized to *Tbp* expression in BAT of FGF21 BKO mice cold exposed for 3 days under *ad libitum*-fed conditions**.**

Data are expressed as means ± SEM. Significant differences were determined by Student's *t*-test, using a significance level of $P < 0.05$.

**Supplemental Figure 2:** mRNA expression of amino acid metabolism genes normalized to *Tbp* expression in BAT of ATF4 BKO mice cold exposed for 4 hours following fasting. Data are expressed as means ± SEM. Significant differences were determined by Student's *t*-test, using a significance level of $P < 0.05$.

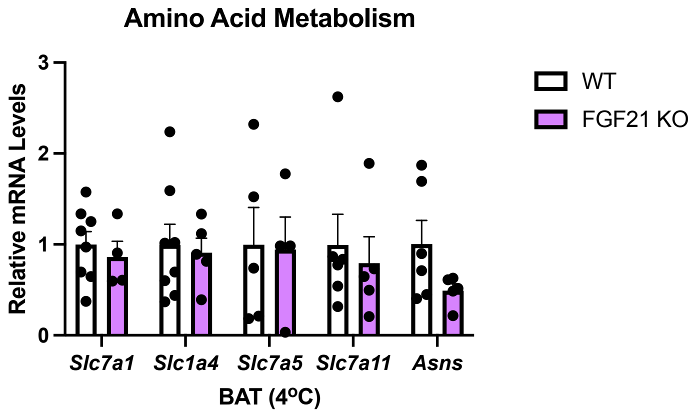


### Supplemental Figure 1

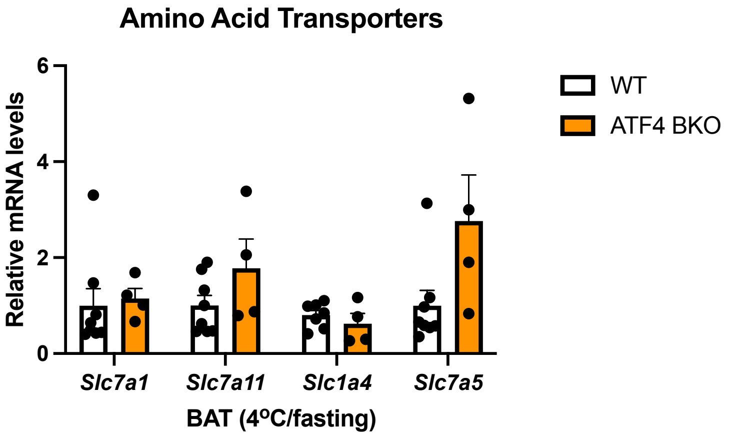


**Supplemental Figure 2**
